# Distinct neurotoxic TDP-43 fibril polymorphs can be generated by heterotypic interactions with α-synuclein

**DOI:** 10.1101/2022.06.14.496041

**Authors:** Shailendra Dhakal, Alicia S. Robang, Nemil Bhatt, Nicha Puangamali, Leiana Fung, Rakez Kayed, Anant K. Paravastu, Vijayaraghavan Rangachari

## Abstract

Amyloid aggregates of specific proteins form important pathological hallmarks in many neurodegenerative diseases, defining neuronal degeneration and disease onset. Recently, increasing numbers of patients show co-morbidities and overlaps between multiple neurodegenerative diseases, presenting distinct phenotypes. Such overlaps are often accompanied by co-localizations of more than one amyloid protein, prompting the question of whether direct interactions between different amyloid proteins could generate heterotypic amyloids. To answer this question, we investigated the effect of α-synuclein (αS) on TDP-43 aggregation inspired by their co-existence in pathologies such as Lewy body dementia and limbic predominant age-related TDP-43 encephalopathy. We previously showed that αS and prion-like C-terminal domain (PrLD) of TDP-43 synergistically interact with one another to generate toxic heterotypic aggregates in vitro. Here, we extend these studies to investigate whether αS induces structurally and functionally distinct polymorphs of PrLD aggregates. Using αS –PrLD heterotypic aggregates generated in two different stoichiometric proportions, we show that αS can effect PrLD fibril forms. The fibril samples have distinctive residue-level structural signatures in NMR spectra, dye-binding capability, proteinase K (PK) stability, and SDS-sensitive thermal stability. By gold nanoparticle labeling and TEM, we show the presence of both αS and PrLD proteins within the same fibrils, and thus the existence of hetertypic hybrid fibrils. We also observe that αS and PrLD co-localize in the cytosol of SH-SY5Y neuroblastoma cells, and show that the heterotypic PrLD fibrils selectively induce synaptic dysfunction in primary cortical neurons. These findings establish the existence of heterotypic amyloid polymorphs and provide a molecular basis for the observed overlap between synucleinopathies and TDP-43 proteinopathies.

## INTRODUCTION

Many neurodegenerative proteinopathies are characterized by the deposition of misfolded protein aggregates (called amyloids) in neuronal and glial cells (1–4). These disorders affect diverse neuroanatomical regions in the brain and exhibit clinical heterogeneity (5–7). Although each protein aggregation disease is often presumed to be caused by misfolding of a single protein, ostensibly distinct neurodegenerative pathologies can show significant clinical and pathological overlap (8–10). It seems as though the phenotype variations may be attributed in part to the interactions between two or more amyloid proteins and accumulation of biochemically distinct heterotypic protein aggregates. For example, aggregates of both tau and αS have been observed in numerous pathologies that are collectively known as tauopathies and synucleinopathies, respectively (11, 12). Indeed, preponderance of interactions between amyloid-β (Aβ), tau, α-synuclein (αS), and transactive response DNA-binding protein 43 kDa (TDP-43) suggests that heterotypic amyloid aggregates may be significant in pathology (13–19). A cornucopia of evidence indicates spatial localization and co-existence of more than one amyloidogenic protein in many proteinopathies. Although not a ubiquitous phenomenon, some amyloid proteins show a greater degree of colocalization than the others such as tau and αS. For example, tau aggregates are often observed in multiple neurodegenerative diseases such as AD, PD, MSA, FTLD etc. (20–22). Similarly, in addition to its presence in Lewy body diseases, aggregates of αS are often observed in the aforementioned pathologies alongside tau deposits (23). Because of their widespread presence in numerous pathologies, the disorders with lesions enriched in tau and αS are termed as tauopathies and synucleinopathies, respectively (21, 24). Yet another protein whose insoluble amyloid inclusions are nearly as widespread as those with tau and αS is TDP-43, and its presence is increasingly becoming known in at least 15 different neurodegenerative diseases including AD, PD, ALS and FTLD, and seems to be a key component in the rapidly expanding spectrum of TDP-43 proteinopathies (25). A sub-set of this spectrum shows overlap with LBDs in which αS aggregates predominate, suggesting colocalization and potential interactions between the TDP-43 and αS. Indeed, mounting evidence indicates synergism between the two protein deposits. In LBD, the severity of αS pathology in the temporal cortex was observed to be significantly more among TDP-43 positive patients, which correlated with the severity of TDP-43 pathology in the amygdala (26). Perhaps the most compelling evidence for selective influence of αS on TDP-43 proteinopathy comes from LATE patient brains in which distinct neuropathological changes are observed in PrLD/αS aggregates containing LATE-LBD and LATE-AD co-pathologies compared to pure LATE (27).

The involvement of TDP-43 in diverse pathophysiological functions has refocused attention to its role in a wide range of neurodegenerative pathologies involving αS aggregates. TDP-43 is a 43 kDa protein that belongs to the ribonucleoprotein family. TDP-43 consists of an N-terminal domain (NTD), two RNA-recognition motifs-RRM1 and RRM2, and a disordered, prion-like C-terminal domain (PrLD) (28, 29). Under physiological conditions, the protein is localized predominantly in the nucleus and plays a role in transcriptional regulation, RNA alternative splicing, miRNA biogenesis, transport, and stabilization (29–31). In pathology, TDP-43 translocates to the cytoplasm, where it undergoes post-translational modifications, including phosphorylation, ubiquitination, and aberrant proteolytic cleavage to generate multiple C-terminal fragments (32–35). These fragments, ranging from approximately 17 kDa (corresponding to PrLD) to 35 kDa, forms insoluble cytoplasmic aggregates and toxic inclusions in cells (33, 36–38). αS is a 14.6 kDa, intrinsically disordered protein containing three domains: amphipathic NTD, central aggregation-prone non-amyloid component (NAC), and an intrinsically disordered charged C-terminal domain (CTD) (39, 40). As noted, although αS and TDP-43 are known to form cytoplasmic amyloid inclusions independently, the question of whether aggregation of the two proteins can be synergistic and coupled to one another has not been addressed in detail. This question is pertinent given the fact that the two proteins are known to influence each other. For example, co-expression of αS and TDP-43 enhances neurodegeneration and loss of dopaminergic neurons in *C.elegans* and transgenic mice (41, 42). Similarly, incubation of exogenous αS fibrils in SH-SY5Y cells enhances TDP-43 phosphorylation and aggregation (43). More compelling pathological evidence of the interaction between the two proteins comes from a recent study which showed that TDP-43 aggregates co-exist with αS aggregates in LATE-LBD and LATE-AD co-pathologies, exhibiting histopathological differences between pure LATE, LATE-LBD and LATE-AD (27). These observations also suggest that the polymorphic fibrils of TDP-43 observed in these patients could be consequential of the influence by αS. Structural polymorphism is known to exist among many amyloid proteins; however, only a few polymorphic strains of TDP-43 fibrils derived from patients have come to the limelight so far (44–48). More importantly, the correlation between structure, biophysical properties and pathophysiology of TDP-43 polymorphs remains unclear. Thus, we ask the question whether polymorphic strains of one amyloid protein can be induced by another amyloidogenic protein. Despite evidence for cross-interactions between amyloid proteins such as those for amyloid-β (Aβ) and islet amyloid polypeptide (IAPP), αS and Aβ, αS and Tau (13, 18, 49–51), the answer to this question remains elusive especially for αS and TDP-43.

Recently, to uncover the mechanistic understanding between the two proteins, we showed that equimolar amounts of αS and PrLD monomers interact synergistically to form co-aggregated, hybrid fibrils (52). In the same report, we also showed that oligomers and fibrils of αS cross-seed PrLD fibril formation selectively, but pre-formed PrLD fibril fails to seed αS monomers hinting at possible conformational selection in cross-interactions between the two proteins. In this report, we extend these investigations to answer the important question of whether cross-seeding of αS to TDP-43 or heterotypic co-aggregation of the two proteins induce specific polymorphic fibrils of TDP-43 compared to the unseeded TDP-43. We show that PrLD aggregation is sensitive to the αS aggregate characteristics; both PrLD–αS hybrid fibrils generated by the interactions of the two monomers and αS fibril (αS^f^)-seeded PrLD fibrils show distinctive differences in structural and biophysical characteristics from PrLD homotypic fibrils formed in the absence of αS. We also show that the two proteins co-localize in the cytoplasm of the SH-SY5Y neuroblastoma cells and that PrLD heterotypic aggregates selectively induce synaptic dysfunction in primary neurons. These results help us understand the generation and propagation of heterotypic αS-TDP-43 amyloid polymorphs and their significance in many neurodegenerative maladies.

## RESULTS

### PrLD and αS co-localize as puncta in the cytoplasm of SH-SY5Y neuroblastoma cells

While co-occurrence of TDP-43 encephalopathy and Lewy body diseases has been reported in the previous studies (13, 27), and our recent study showed synergistic interactions between PrLD and αS (52), it is unclear whether the two proteins co-localize in the cytoplasm. To investigate this, confluent SH-SY5Y neuroblastoma cells were separately transfected with the plasmids of blue fluorescent protein (sBFP2)-tagged PrLD and sBFP2 alone as a control. After 24 hours of incubation at 37°C to allow transient expression, both the PrLD-expressing and control cells were pulsed with Hilyte-532-labeled 500 nM recombinant αS monomers prepared in 20 mM MES buffer (detailed in Methods section) and incubated for additional 24 hours to ensure the maximum internalization as reported previously (53). Live-cell imaging after adding nuclear stain under a confocal microscope revealed that the control sBFP2 expressing cells exhibited diffused blue fluorescence throughout the cells with a few denser foci in the cytoplasm. At the same time, the internalized αS monomers were present in the diffused form in the cytoplasm as anticipated (Figure 1a). Thus, sBFP2 and αS did not show colocalization but remained primarily diffused. In contrast, sBFP2-PrLD expressing cells showed distinct fluorescent puncta present exclusively in the cytoplasm, which we believe are PrLD aggregates (Figure 1b). Interestingly, internalized αS (yellow) showed colocalization within the puncta of PrLD (Figure 1b). Confirming cytoplasmic colocalization of the two proteins provides a cellular basis and support for their observed direct interactions (49) and potential formation of pathological polymorphs.

**Figure 1.**
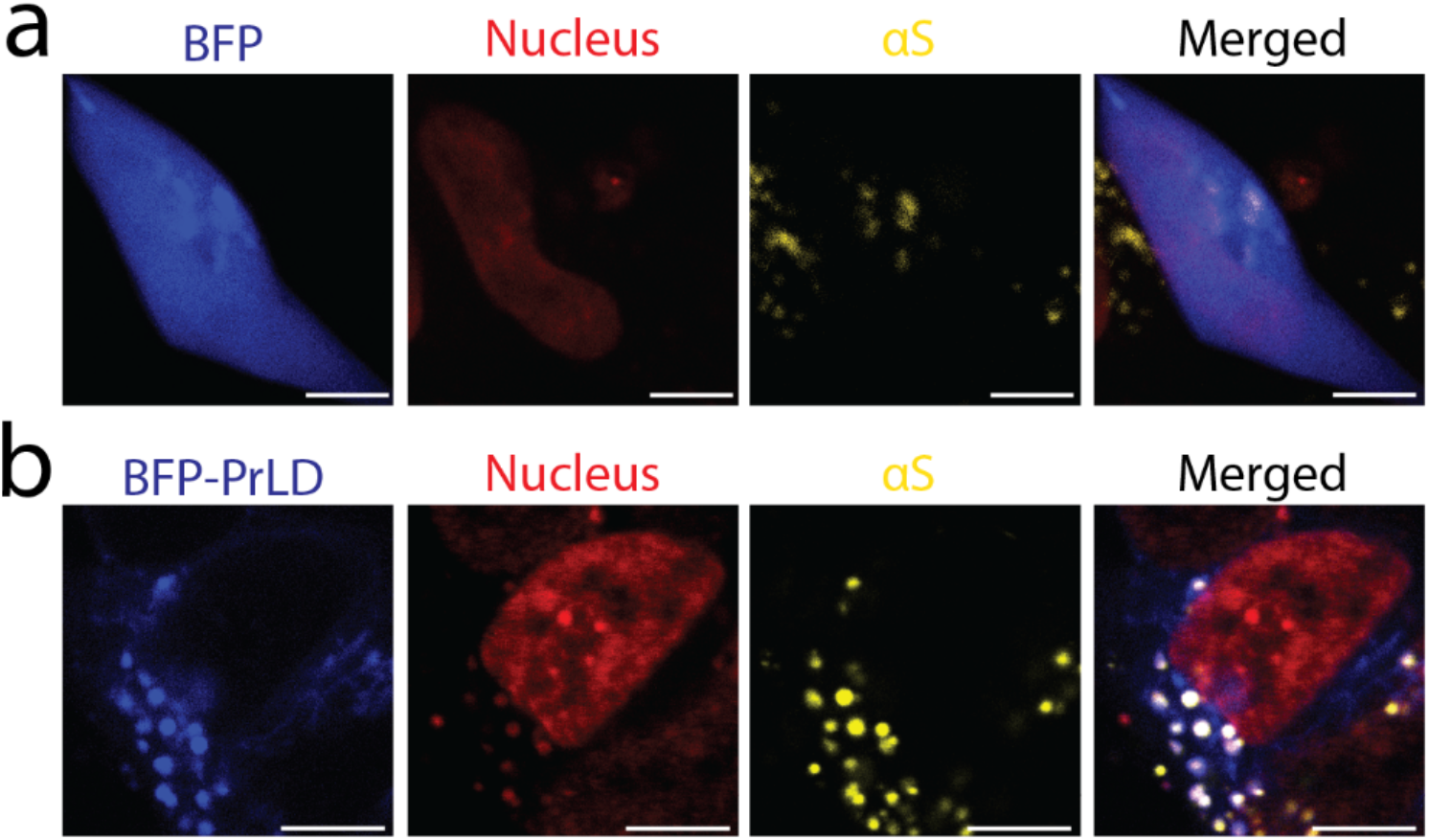
Representative live-cell images showing colocalization of αS with blue fluorescent protein (BFP) and BFP tagged prion-like domain (BFP-PrLD). Fluorescent confocal images of human SH-SY5Y neuroblastoma cells 48 hours after transfection with BFP (a) and BFP-PrLD (b). Cells were pulse-chased with Hilyte-532 fluorescently labeled monomeric recombinant αS (yellow) 24 hours post-transfection. Cells were stained with nuclear stain (red) prior to confocal live-cell imaging at 40x magnification. Merged panel shows degree of cytoplasmic colocalization of αS with BFP and BFP-PrLD. (Scale bar = 5 μm).

### Gold nanoparticle-decorated transmission electron microscopy (TEM) images support the existance of heterotypic hybrid fibrils of αS and PrLD

First, morphological features of heterotypic aggregates were investigated by negative staining TEM. Both homotypic PrLD and αS fibrils showed smooth and long unbranched fibrils as expected (Figure 2a and b). In contrast, both PrLD–αS hybrid and αS^f^-seeded PrLD fibrils showed more branching and clumping in their fibrillar structures (Figure 2c and d). To quantify the amounts of αS and PrLD within the fibrils, samples were treated with formic acid (to disaggregate the fibrils), and subjected to MALDI-ToF analysis. The homotypic PrLD showed the expected mass (Figure 2e). The PrLD–αS hybrid fibrils showed the presence of approximately equimolar amounts of αS and PrLD as shown in our previous study (52) (Figure 2f). Since αS fibrils were added as sub-stoichiometric seed, as expected, αS was barely detectable in the αS^f^-seeded PrLD fibrils (Figure 2g). All the samples also showed low amounts of sinapinic acid adducts, which has been shown to be common on this matrix (54). To establish the formation of heterotypic aggregates, PrLD–αS hybrid fibrils were investigated using Au-nanoparticle -labeled TEM. Sub-stoichiometric hexa-histidine tagged protein to facilitate nanogold particle binding was mixed with the untagged protein (hetero or homo) and incubated for fibril formation (see Methods). Fibrils were then isolated and incubated with Au-nanoparticles to be visualized by TEM (Figure 2h). The positive control, homotypic PrLD fibrils were prepared by mixing 5 μM his-tagged PrLD with 20 μM untagged PrLD fibrillar structures studded with Au-nanoparticles (Figure 2i). Images of Au-nanoparticle incubated PrLD–αS hybrid fibrils containing histagged αS and untagged PrLD also showed the presence of Au-nanoparticle studded fibrils confirming the presence of both PrLD and αS within the sample fibrils. In other words, there is a hybrid heterotypic PrLD–αS polymorph (Figure 2j).

**Figure 2.**
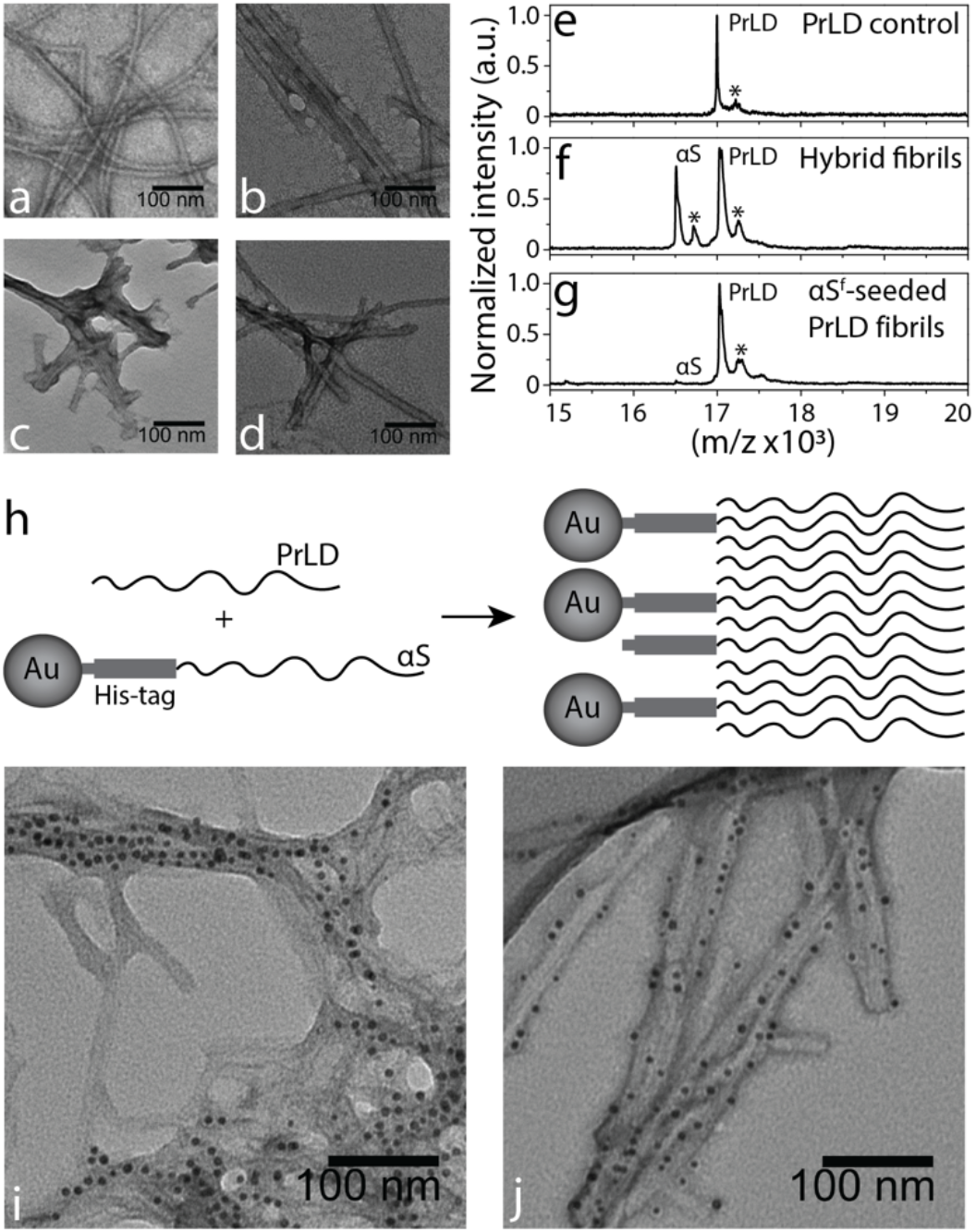
TEM images of unlabeled and nanogold-labeled fibrils along with their MALDI-ToF spectra. a-e) Negative EM staining of (a) homotypic PrLD fibrils, (b) homotypic αS fibrils (c) hybrid fibrils, and (d) αS^f^-seeded PrLD fibrils. e-g) Normalized MALDI-ToF spectra of formic acid treated PrLD homotypic fibrils (e), hybrid fibrils (f), and αS^f^-seeded PrLD fibrils (g) showing relative amount of αS and PrLD monomers. The ‘*’ indicates the protein adduct formed with sinapinic acid (SA). h) Schematic showing possible arrangement of Au-nanoparticle bound his-tagged αS with untagged PrLD in the hybrid fibrils. i-j) Ni-NTA Au-nanoparticle-labeled PrLD fibrils (i), and hybrid fibrils (j). Black dots on the fibrils are 5-micron nanogold particles bound to the hexa-histidine tag on αS or PrLD surface.

### αS-induced PrLD fibril polymorphs exhibit biophysical differences

To further examine the conformational differences among PrLD polymorphs generated by the interactions with αS, intrinsic tryptophan fluorescence along with binding to known amyloid dyes such as 8-anilinonaphthalene-1-sulfonic acid (ANS), curcumin and 9-(dicyano-vinyl) julolidine (DCVJ) were measured. First, homotypic PrLD fibrils generated in the absence of αS, hybrid fibrils generated by incubating equimolar amount of PrLD and αS monomers and αS^f^-seeded PrLD fibrils along with controls were analyzed for intrinsic tryptophan fluorescence. Fibrils were prepared as reported in our previous study (52), (detailed in Methods section). All three samples, PrLD–αS hybrid fibrils, αS^f^-seeded PrLD fibrils and homotypic PrLD control fibrils showed blue shifts compared to PrLD monomers (λ^Em^ = 340 nm), reflecting a solvent-protected apolar environment within the fibrils (Figure 3a). Among the samples, hybrid fibrils showed marginally greater blue shift (Δl = |12| nm) as compared to αS^f^-seeded PrLD (Δl = |8| nm) or control PrLD fibrils (Δl = |6| nm) (Figure 3b).

**Figure 3.**
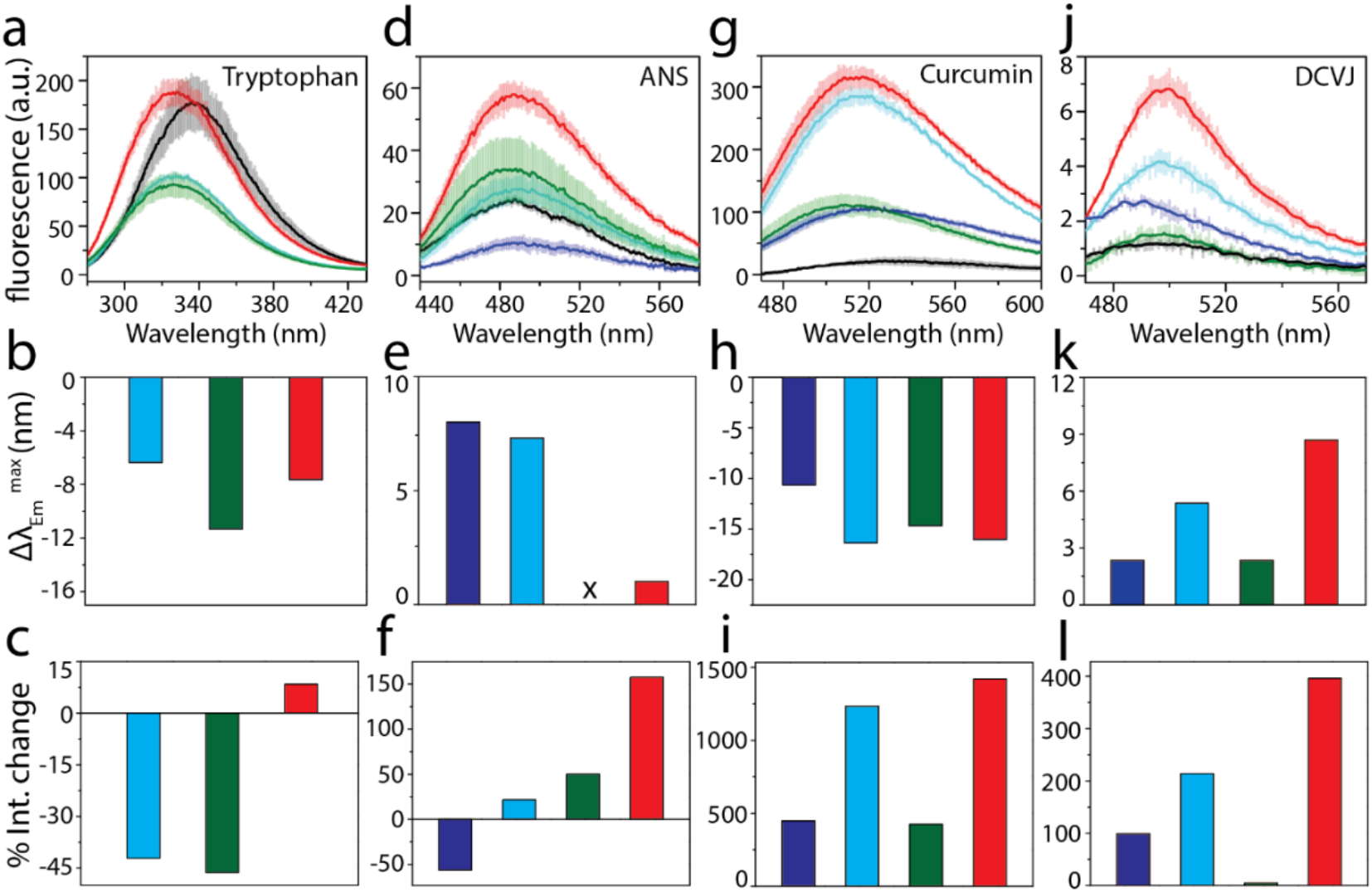
Intrinsic fluorescence and dye binding of monomeric and fibrillar αS or TDP-43 PrLD species. Intrinsic tryptophan fluorescence emission scans of monomeric TDP-43 PrLD, TDP-43 PrLD fibrils, αS-TDP-43 PrLD hybrid fibrils, and αS^f^-seeded TDP-43 PrLD fibrils (a). Mean shift in tryptophan emission maxima (b) and percentage change in total intensity (c) of TDP-43 PrLD fibrils, αS-TDP-43 PrLD hybrid fibrils, and αS^f^-seeded TDP-43 PrLD fibrils to TDP-43 PrLD monomers. d-l) Fluorescence emission scans for ANS (d), curcumin (g) and DCVJ (j) dyes. Mean shift in emission maxima (Δλ_Em_^max^) and percentage change in total intensity (% Int. change) of αS monomers, αS and TDP-43 homotypic fibrils, αS-TDP-43 PrLD hybrid fibrils, and αS^f^-seeded TDP-43 PrLD fibrils to TDP-43 PrLD monomers in ANS (e-f), curcumin (h-i) and DCVJ (k-l) dyes. Both monomers and fibrils (2 μM) were prepared in 20 mM MES buffer pH 6.0. Samples concentration was determined using Pierce® BCA protein assay kit; fibrils concentration is expressed as monomer equivalents (see details in materials and method section). αS monomers (—), TDP-43 PrLD monomers (—), αS fibrils (—), TDP-43 PrLD fibrils (—), hybrid fibrils (—), and αS^f^-seeded TDP-43 PrLD fibrils (—) are indicated using the respective colors.

However, the intensities for both control PrLD and PrLD–αS hybrid fibrils showed a significant decrease while αS^f^-seeded PrLD fibrils showed a small increase in intensity reflecting a more solvent-protected environment for latter (Figure 3c). Intrinsic fluorescence for control αS fibrils could not be monitored as the protein is devoid of tryptophan residues. The ANS dye, known to binding exposed hydrophobic surfaces (55, 56), also showed significant differences in both wavelength shifts and intensities (Figure 3d-f).

Surprisingly, PrLD–αS hybrid fibrils showed no wavelength shift compared to monomers but both control PrLD, and αS^f^-seeded PrLD fibrils showed red shifts in the order of (Δλ = |7| nm) and (Δλ = |2| nm), respectively (Figure 2e). The intensity of ANS fluorescence, which reflects the extent of exposed hydrophobic surfaces, showed noticeable changes (Figure 3f); here too, αS^f^-seeded PrLD fibrils showed the largest % increase, followed by PrLD–αS hybrid fibrils and the control PrLD fibrils, while the control αS^f^ showed a decrease in the intensity (Figure 3f). Curcumin which is known to bind and distinguish different aggregate structures (57, 58), also showed significant blue shifts in wavelength that were somewhat uniform in magnitude (Δλ = |14| nm) except for control αS fibrils (Δλ = ~|10| nm) (Figure 3g and h). The intensities for the same samples, however, showed variations; again, αS^f^-seeded PrLD fibrils showed the largest % increase (1500%), followed by control PrLD fibrils (1100%), while PrLD–αS hybrid fibrils and control αS fibrils showed lesser change (500 %) (Figure 3i). Finally, the addition of the dye DCVJ showed dramatic redshifts in wavelength for αS^f^-seeded PrLD fibrils (Δλ = |9| nm) followed by control PrLD (Δλ = |5.5| nm), PrLD–αS hybrid (Δλ = |2.5| nm) and control αS^f^ (Δλ = |2.5| nm) (Figure 3j and k). The αS^f^-seeded PrLD fibril sample also showed a dramatic % increase (390%) and PrLD–αS hybrid fibrils showed no change in the intensity, and control PrLD and αS fibrils showed 200 and 100% increases, respectively (Figure 3l). Together, these data bring out the subtle yet important differences in the ability of the amyloid binding dyes to distinguish between fibril polymorphs of PrLD induced by αS. More importantly each of the dyes were able to differentiate between PrLD–αS hybrid, αS^f^-seeded, and control PrLD fibrils, bringing out the potential conformational differences between these three polymorphs.

To investigate the potential differences between PrLD–αS hybrid, αS^f^-seeded PrLD, and control PrLD fibril polymorphs, their enzymatic stabilities were analyzed. First, to see how stable the polymorphs are towards enzymatic degradation, a broad-spectrum serine protease-proteinase K (PK), that is widely used to assess the stability and conformational differences of many amyloid proteins was used (59–61). The three fibril samples along with control αS fibrils were incubated with PK, and aliquots of the samples were quenched at various time points to investigate the kinetic stability of the fibrils for degradation. The samples were then analyzed using SDS-PAGE and MALDI-ToF mass spectrometry. PK digestion of control αS^f^ showed a predominant band at ~28 kDa (Figure 4a) but without other low molecular weight bands, presumably due to extensive digestion. Control PrLD fibrils showed bands near ~10, 17, and 30 kDa along with a faint band smear, presumably due to the dissociation of higher molecular weight oligomers and fibrils (Figure 4a). In addition to the predominant bands near 28 and 10 kDa, PrLD–αS hybrid fibrils and αS^f^-seeded PrLD fibrils showed multiple digested bands near 50, 28-10, and 6 kDa that are different from those observed for the control fibrils (arrows; Figure 4a). The temporal stability of the polymorphic fibrils also showed differences in the rate of digestion.

**Figure 4.**
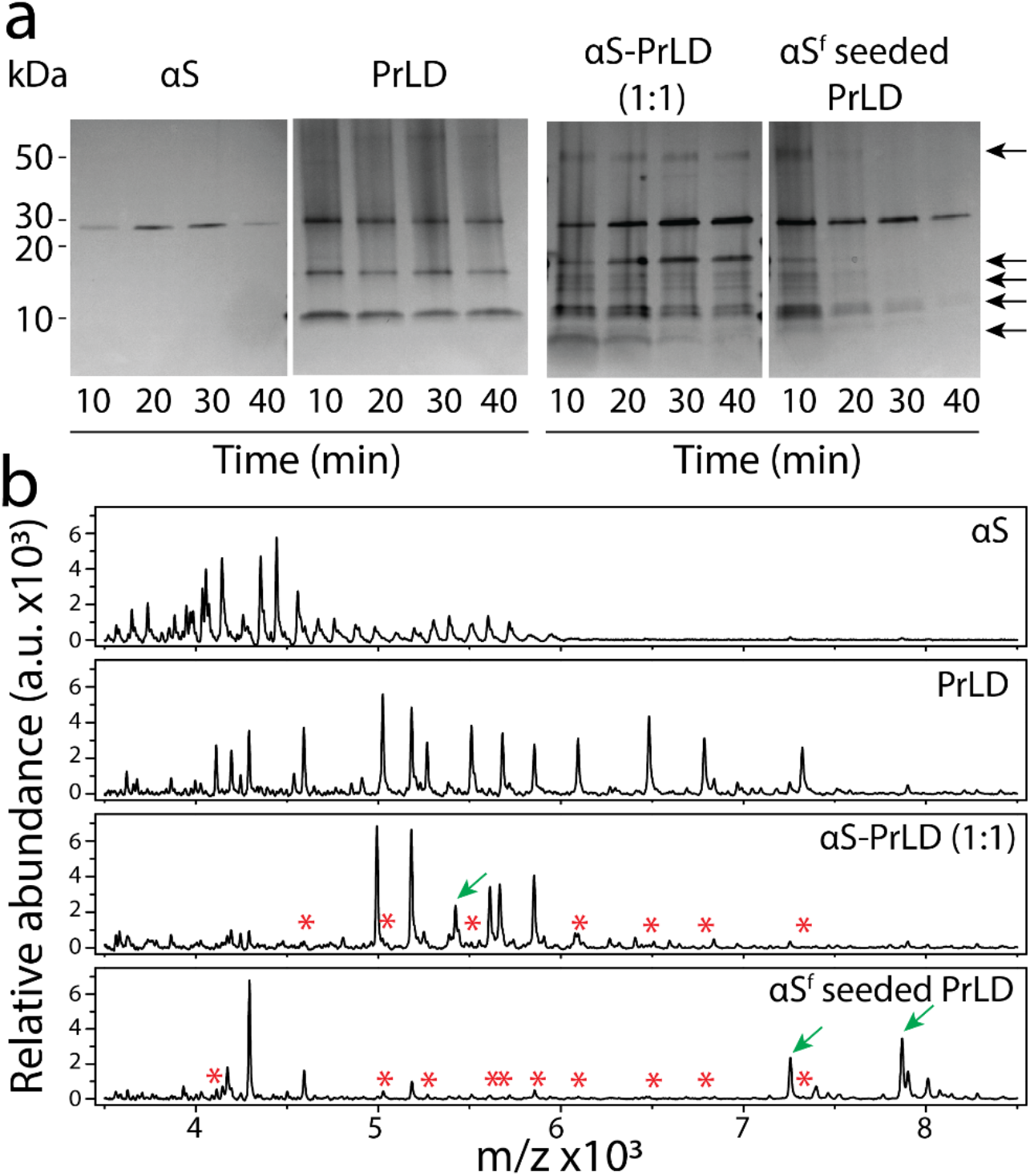
SDS-PAGE gel and MALDI-ToF spectra of Proteinase K digested polymorphic αS and PrLD fibrils. a) Proteinase K digestion of αS homotypic fibrils, PrLD homotypic fibrils, αS-PrLD (1:1) hybrid fibrils and αS^f^-seeded PrLD fibrils at different time points (10, 20, 30 and 40 minutes). Arrows indicate the digested bands unique to heterotypic fibrils. b) MALDI-ToF of proteinase K digested fragments from the same sample. Arrows (→) indicate the fragments unique to heterotypic fibrils, and star (*) indicate fragments that are absent in heterotypic ones.

While the control fibrils showed barely any change in the intensity of the digested bands, the bands near 8, 10, and 50 kDa for the PrLD–αS hybrid and αS^f^-seeded fibrils showed progressive disappearance with time, suggesting lesser enzymatic stabilities for the two fibrils (Figure 4a). This data also indicates that heterotypic fibril structures have more accessibility for PK to cleave the polypeptide backbone in core-amyloid regions or in other words, more amorphous than homotypic fibrils. To investigate the digestion pattern in the low molecular weight range (< 8 kDa) that is not visible in SDS-PAGE gels, the same samples were analyzed by MALDI-TOF spectrometry. Homotypic control αS fibrils showed extensive digestion with numerous small molecular fragments (< 6 kDa), which also answers why higher molecular weight fragmentations were not observed in the gel (Figure 4b). The control PrLD fibrils also showed many fragments between 5 and 7 kDa (Figure 4b). In contrast, the heterotypic fibril polymorphs showed fewer digested fragments but, more importantly, showed the absence of the digested fragments observed in the control along with some newer fragments suggesting a different structure for all the polymorphs (Figure 4b).

Based on the PK digestion patterns, one could argue that the heterotypic fibril polymorphs of PrLD–αS hybrid, αS^f^-seeded fibrils are more exposed to PK than the control homotypic PrLD fibrils. Therefore, it can be conjectured that the heterotypic polymorphs may show lesser thermodynamic stabilities than the homotypic PrLD fibrils. To test this interpretation, all samples were subjected to SDS treatment and denaturation and disaggregation as function of temperature. As established previously in our lab, SDS pretreatment can help reveal relative stabilities of amyloid aggregates since direct temperature increase does not “melt” amyloid aggregates (58). Typically, the conversion of β-sheets (fibrils) to α-helical structures (SDS-denatured and disaggregated) is monitored by far-UV CD as a function of temperature, and the degree of melting is represented as the ellipticity difference between 208 nm (α-helix) and 218 nm (β-sheet); more positive the difference is, more β-sheet the structure is.

The control homotypic PrLD fibrils, prior to SDS treatment and temperature increase, displayed an intense β-sheet spectrum with a minimum at 218 nm (pre-melt; Figure 5a). An increase in temperature resulted in a conformational change to an α-helical structure with a mid-point of transition at ~60 °C, typical for an amyloid fiber (Figures 5a and e). The fibrils also failed to convert to a complete α-helix even after 90 °C, suggesting high thermal stability (Figure 5e). Heterotypic PrLD–αS hybrid fibrils showed a much less intense spectrum with a broad minimum at 214-218 nm suggesting a less well-defined β-sheet signature prior to melting (pre-melt; Figure 5b). Treatment of SDS immediately converted the spectrum to a partial α-helical structure which became more defined and more intense with the increase in temperature (Figures 5b and e). Caution must be exercised with the intensity of the spectra as the data were normalized based on our assumed 1:1 stoichiometry for PrLD: αS within the fibrils (Figure 1a) (52). Nevertheless, nearly half of the β-sheet melted to helix at 20 °C, and completely at 40 °C, which suggests that the fibrils are more amorphous and less stable than the homotypic PrLD ones (Figure 5e). αS^f^-seeded PrLD fibrils showed an intense minimum at 218 nm (β-sheet) with a shoulder near 210 nm before melt, indicating the presence of an α-helical component within the fibrils to some degree (pre-melt; Figure 5c). This result suggests a structure that is different from the homotypic fibrils. Treatment with SDS at 20 °C showed nearly half of the structure converted to an α-helix similar to PrLD–αS hybrid fibrils (Figure 5c and e). But subsequent melting upon increasing temperature showed a slow and gradual conversion to a complete helical structure at 90 °C (Figure 5e). As expected, the control PrLD monomers showed a predominant random coil structure with a partial α-helix as it is known to have a helical segment in the middle (62) that almost immediately melted to a complete α-helix (Figure 5d and e). The date collectively indicates that both PrLD–αS hybrid and αS^f^-seeded PrLD fibrils differ in their structures, sensitivity towards SDS denaturation and thermal stability from homotypic PrLD fibrils. Both heterotypic fibrils seem to be more amorphous, SDS-unstable fibrils, which seems to correlate with the PK enzymatic stability.

**Figure 5.**
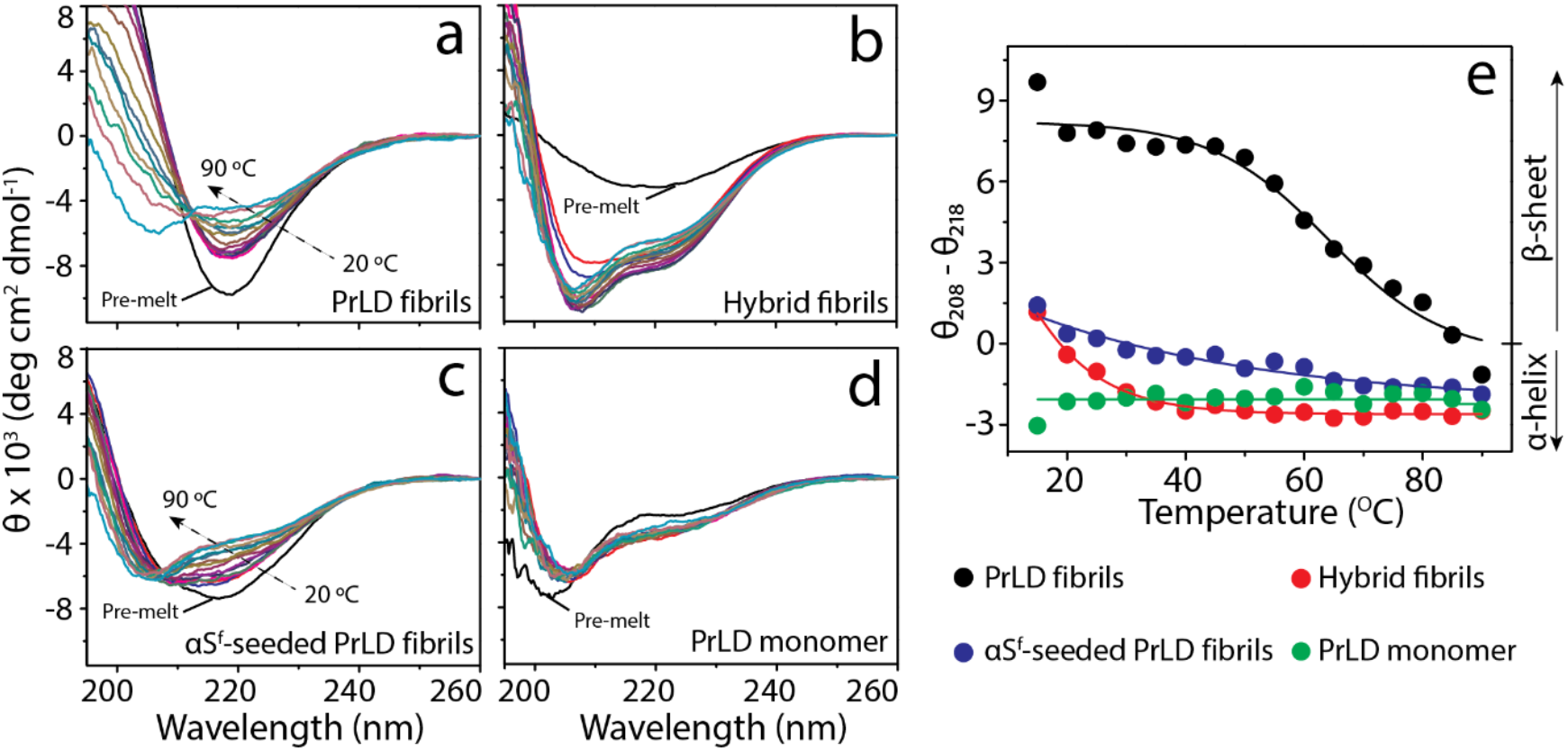
Differences in the thermal stabilities of αS-induced PrLD polymorphs suggest differences in their core structures. a-d) Mean residue ellipticity [θ] from circular dichroism (CD) spectra obtained for homotypic PrLD fibrils (a), PrLD–αS hybrid fibrils (b), αS^f^-seeded PrLD fibrils (c) and PrLD monomers (d) after treatment with 1% SDS and temperature increase from 20 °C to 90 °C. e) The difference in [θ] between 208 nm (α-helix) and 218 nm (β-sheet) is plotted as a function of temperature and fitted with Boltzmann’s sigmoidal fits.

### PrLD amyloid molecular structure is sensitive to αS interactions

NMR spectra indicate that PrLD molecular structure differs between PrLD–αS hybrid, αS^f^-seeded PrLD fibrils, and control PrLD control fibrils (Figure 6). In these experiments, only PrLD was isotopically labeled with ^13^C. The spectra directly report on PrLD structure influenced by the interactions with αS. The molecular structure of PrLD within the amyloid core depends on whether PrLD fibrils are homotypic, formed in the absecne of αS (Figure 6a), αS^f^-seeded PrLD fibrils (Figure 6b), or PrLD–αS hybrid fibrils (formed by equimolar co-incubation of PrLD and αS monomers in solution) (Figure 6c). The NMR techniques employed (cross polarization as well as DARR dipolar recoupling (63, 64)) are expected to yield signals from rigid regions of the spectrum (65), and thus produce signals only from the amyloid core structure. Furthermore, for the 50 ms mixing time employed for ^13^C-^13^C dipolar couplings the off-diagonal peaks are mainly due to ^13^C-^13^C interactions between atoms within the same amino acids or adjacent amino acids in the primary structure. Noting that the primary structure of the PrLD or the NMR experimental parameters did not vary between samples, the appearance of NMR crosspeaks in some spectra and not others indicates sample-dependent differences in the amyloid core regions. It is expected that protein backbone motion in the presence of water (samples were hydrated ultracentrifuge pellets) would suppress crosspeaks for molecular domains outside of the amyloid core. Another factor that would reduce detectability of signals outside the amyloid core is inhomogeneous broadening, caused a greater distribution of molecular conformations expected for regions outside of the amyloid core. To further illustrate the differences between the samples, Figure 6d shows overlaid ^13^C spectra for the three fibril samples.

**Figure 6.**
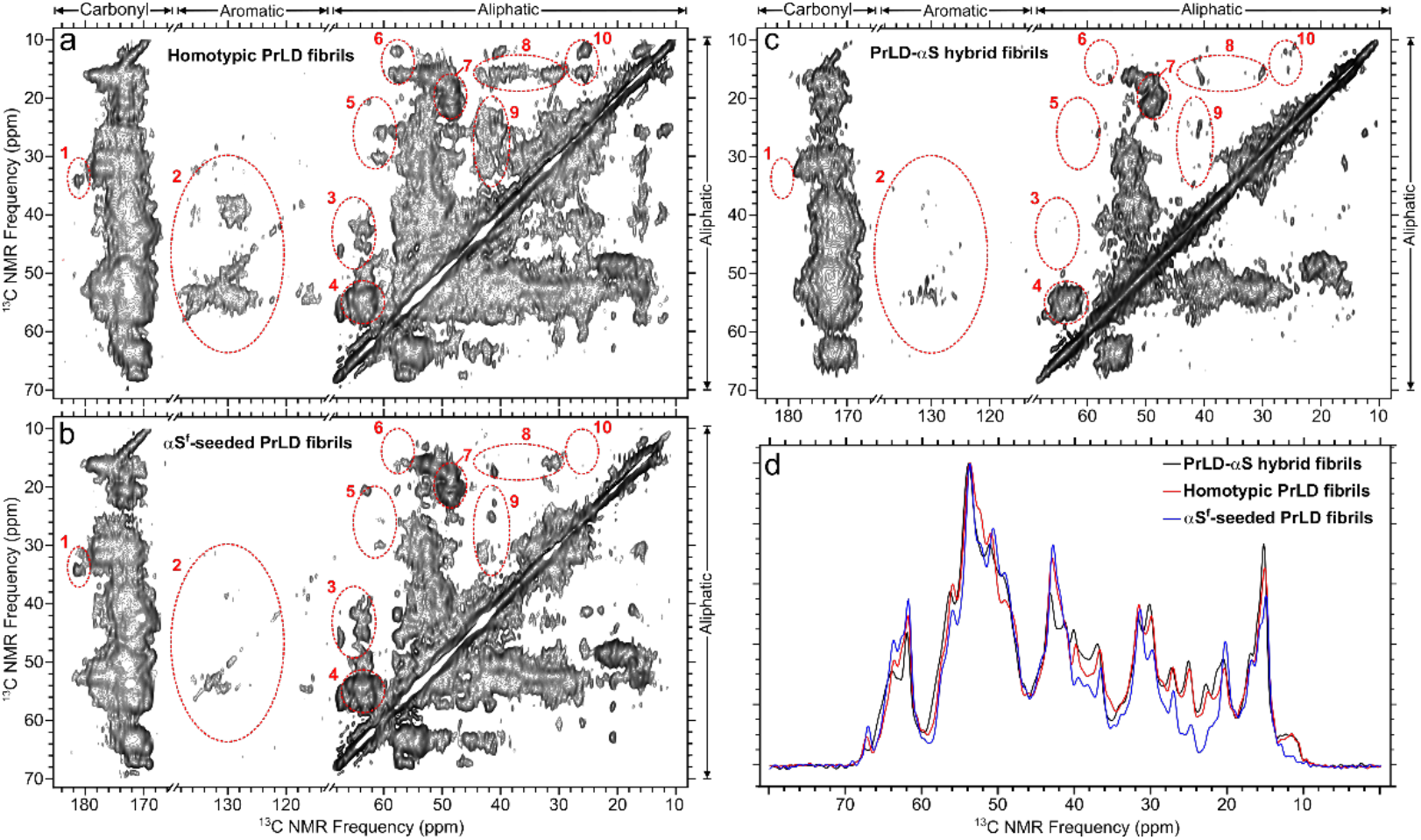
^13^C NMR spectra of uniformly ^13^C-labeled TDP-43 PrLD in amyloid fibrils. Spectra correspond to: 2D ^13^C-^13^C unseeded homotypic PrLD fibrils (a), αS^f^-seeded PrLD fibrils (b), and heterotypic PrLD–αS hybrid fibrils (c) and overlaid aliphatic regions (scaled for equal total intensity) of the ^13^C NMR spectra from all three samples (d). The αS protein was not labeled with ^13^C and therefore did not contribute directly to the spectra. The regions of (a-c) marked by red dotted ellipses indicate spectral features that differ between spectra, as discussed in the text.

Although further research is necessary to determine the detailed effects of αS on TDP-43 aggregated structure, molecular structural variation is clearly evident in Figure 6. To guide the eye, we circled selected regions of the spectrum, and further note that some residues occur sparsely in the PrLD seqeunce. The regions 1, 2, 6, 8, 9, and 10 in Figure 6 indicate crosspeaks between ^13^C atoms in residues that are low-abundance or rarely occur in the PrLD sequence (including E, D, Y, W, I, and L) (66). These regions also include several crosspeaks that appear in some but not all the PrLD spectra. The regions 3 and 5 include crosspeaks between atoms on nearest-neighbor amino acids, most likely residues adjacent to S residues. Regions 4 and 7 correspond to the high-abundance amino acids S and A. The homotypic PrLD fibrils may have a structure like the cryo-EM structure reported earlier (67), in terms of residues involved in the core fibril. In contrast, we detected fewer signals overall in the spectra from αS-seeded and heterotypic fibrils (Figures 6b and 6c, respectively), suggesting that these structures include smaller (or entirely distinct) regions of PrLD.

### αS^f^-seeded PrLD fibrils cause synaptic dysfunction in primary neuronal cultures

To determine the functional consequences of αS-induced TDP-43 polymorphs, we investigated their effect on synapses. C57BL/6 primary cortical neurons were treated with 1μM of αS^f^-seeded PrLD fibrils along with homotypic αS and PrLD fibrils. The monomers of each protein were also used as controls. The cells were fixed after 24h and immunostained using pre-(Synapsin 1) and post-synaptic (PSD95) antibodies (Figure 7a). The cells were then imaged using a confocal microscope and analyzed using the protocols described in the methods section. The αS^f^-seeded PrLD fibrils (*p* = 0.0138), homotypic αS and PrLD fibrils (*p* = 0.0013) and αS monomers (*p* = 0.0089) showed reduction in Synapsin 1 density when compared to the untreated cells (Figure 7b). But the reductions were ~ 10-15% as compared to the untreated cells (Figure 7b). In contrast, a significant reduction in PSD95 density was observed when the neurons were treated with αS^f^-seeded PrLD fibrils (~ 35%; *p* < 0.001), followed by PrLD and αS monomers (~18-20%) as compared to the untreated control (Figure 7c). Analysis of the colocalization of the pre- and post-synaptic signal density also showed that αS^f^-seeded PrLD fibrils treated neurons had the greatest reduction compared to untreated cells (~ 50%; Figure 7c). This suggests that selectively, αS^f^-seeded PrLD fibril polymorph is able to reduce the number of functional synapses in primary neurons. In sum, the data indicate that αS^f^-seeded PrLD fibril polymorph could contribute to synaptic dysfunction.

**Figure 7.**
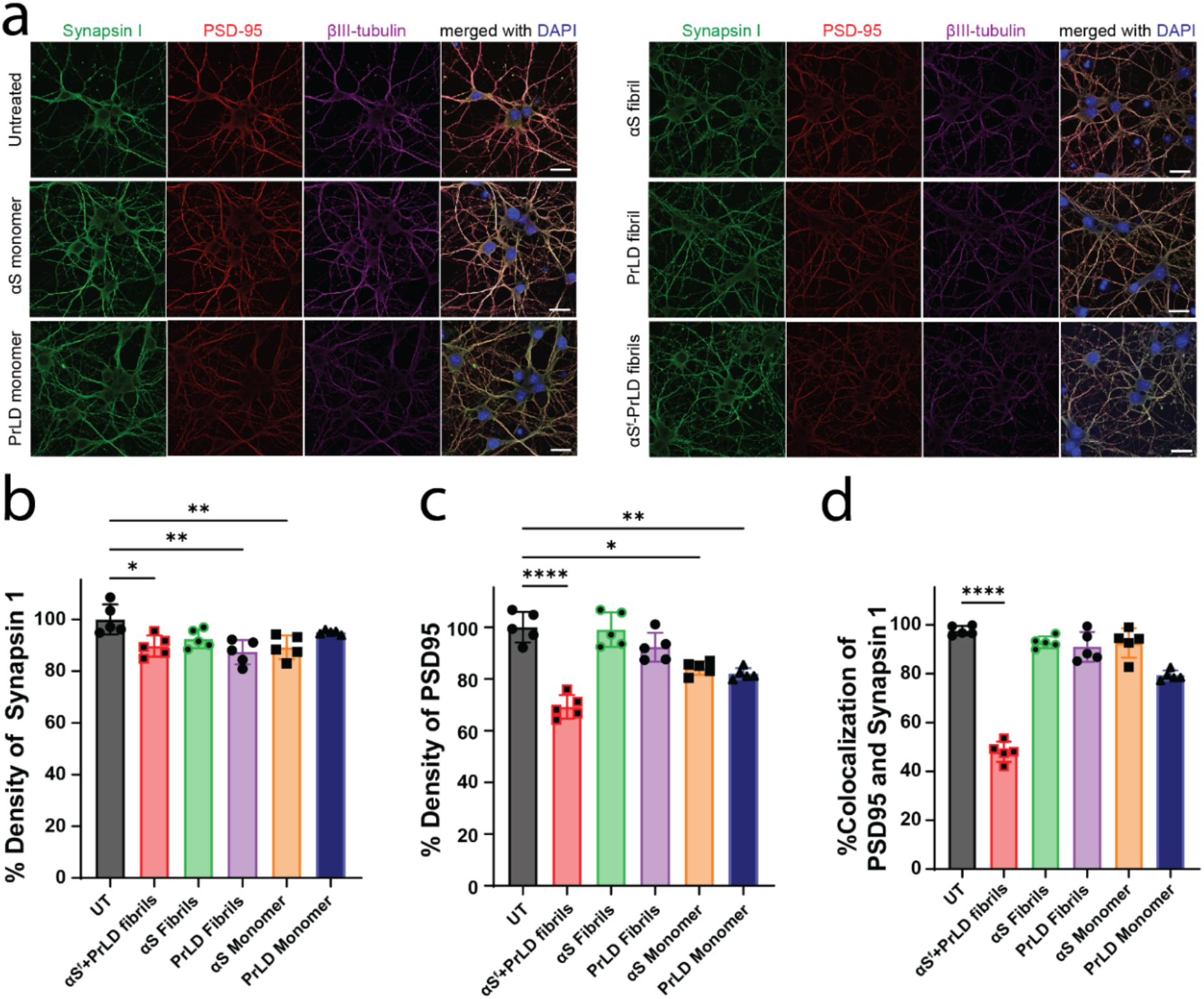
Effect of monomeric and fibrillar αS and TDP-43 PrLD along with αS^f^-seeded TDP-43 PrLD fibrils on pre- and post-synaptic expression in primary cortical neurons. (**a**) Representative confocal images of C57BL/6 primary cortical neurons treated with 1μM monomer (left panel) or fibril (right panel) of α-Synuclein (αS), prion-like domain (PrLD) of TDP-43, or αS fibrils seeded TDP-43 PrLD fibrils (αS^f^-PrLD) for 24 h. Cells were fixed and immunostained with pre-synaptic marker (Synapsin I, green), post-synaptic maker (PSD-95, red), mature neuronal marker (βIII-tubulin, magenta), and merged with nuclear staining (DAPI, blue). Scale bar = 20 μm. (**b-d**) Quantification of density of pre-synaptic (**b**), post-synaptic (**c**), and Pearson’s colocalization coefficient analysis of the synaptic proteins (**d**) in αS, PrLD, or αS^f^-PrLD-treated primary neurons using Imaris software. Each treatment group was randomly imaged in five different regions of interest and performed in triplicate. Image analysis was calculated by one-way ANOVA with Tukey’s multiple comparison. Bar graphs represent the value of mean ± SD. \**p* < 0.05, \*\**p* < 0.01, \*\*\*\**p* < 0.0001.

## DISCUSSION

Significant clinical and pathological overlaps observed among neurodegenerative diseases have brought to bear increased attention on understanding the potential cross-interactions between different amyloid proteins. Many amyloid proteins are known to cross-interact with one another both in vitro and in vivo including Aβ and hIAPP (68–70), αS and tau (71, 72), αS and Aβ (73, 74). Recently, tau was shown to induce distinct molecular conformations of αS filaments in vitro, implicating that heterotypic interactions may generate specific polymorphs and potentially discrete phenotypes (75). However, the involvement of heterotypic aggregates formed as a result of co-aggregation of two different proteins to behave as polymorphic strains remains unclear. Compounding this scarcity is the lack of high-resolution structures of brain-derived TDP-43 polymorphs. In fact, only a few examples exist that include the cryo-EM structure of TDP-43 PrLD fibrils. For example, TDP-43 fibrils derived from frontal cortex of an ALS patient showed a unique ‘double-spiral fold’ (76). This was different from the cryo-EM structure of PrLD fibrils generated in vitro (67). In addition, two different amyloid forming segments of TDP-43, SegA (residue 311-360) and SegB A315E (386-331), revealed four different polymorphs (44). TDP-43 strains have also been isolated from FTLD-TDP brain tissues, which induces the formation of morphologically distinct aggregates in cells, and induces distinct morphological and subcellular distribution of TDP-43 pathology in transgenic mice (45). Although these studies showed TDP-43 polymorphs both in vitro and in vivo, TDP-43 strains in co-pathologies are not investigated so far. In our previous study, we discovered that αS monomers, oligomers, and fibrils are able to promote PrLD fibrillization (52), further providing a possible molecular mechanism by which the two proteins may interact with one another. This current report advances these findings by showing that the two proteins indeed form co-localized cytoplasmic foci in SH-SY5Y neuroblastoma cells. More importantly, the data unequivocally show that depending on the nature of the αS seeds, i.e, whether they are monomers or fibrils, conformationally distinct polymorphs of PrLD are generated. However, it is interesting to observe that addition of αS (monomers or fibrils) seems to destabilize heterotypic PrLD fibril polymorphs by making them more amorphous (evident from PK digestion and temperature stability data). Equimolar incubation of both monomers generates fibrils in which the proteins retain their equimolar composition suggesting ‘hybrid’ fibrils (52). Au-nanoparticle labeling and TEM of PrLD–αS hybrid fibrils shown here unequivocally suggests integration of αS within the PrLD fibrils. Although it is clear that hybrid fibrils are less stable and more susceptible towards enzymatic digestion, they show residue-level differences in solid-state NMR suggesting a different structure as compared to either αS^f^-seeded or unseeded PrLD. Despite being less stable than homotypic PrLD fibrils, the αS^f^-seeded PrLD fibrils show significantly different synaptic effect in primary neurons suggesting amorphous polymorph may functionally be more deleterious in vivo.

It is important to distinguish the differences between the cross-seeding and heterotypic co-aggregation. As discussed throughout the paper, several examples of cross-seeding interactions among amyloid proteins exist. Typically, cross-seeding occurs when a pre-formed seed of one amyloid protein enhances the aggregation of a second amyloid protein. Such a cross-seeding may result in fibrils that are similar or dissimilar to homotypic fibrils without the influence of the second amyloid protein seed. On the other hand, co-aggregation is a process of synergistic aggregation of two different amyloid protein monomers towards heterotypic hybrid fibrils containing both proteins. Hybrid heterotypic aggregates thus far have not been observed among amyloid proteins except for a select few atypical amyloid systems (77–80). PrLD–αS hybrid fibrils shown here and in our previous study (52) form the first among the class of hybrid heterotypic amyloid fibrils. Furthermore, the results presented here suggest that both heterotypic PrLD fibrils (cross-seeded or co-aggregated with αS) have different structures and biophysical properties.

Among the two modes of cross-interactions of PrLD with αS, i.e, αS monomers or αS^f^, the latter is more pathologically relevant as pre-existing Lewy bodies can seed TDP-43 aggregates in the cytoplasm. Although, reverse phenomenon involving PrLD^f^ seeding αS monomers could also be equally relevant and interesting, this was not investigated because our earlier data demonstrated that seeding of αS^f^ to PrLD monomers is far more efficient than PrLD^f^ seeding αS monomers suggesting conformational selectivity in seeding (81–83). The data presented here establish that aggregation dynamics and structures of PrLD fibrils grown under the influence of αS are different from the homotypic PrLD fibrils. Importantly, polymorphism seems to be correlate with pathological changes in primary neurons. However, several questions still remain less understood. For example, how PrLD fibril polymorph generated via templated seeding of αS^f^ is different from those generated by co-aggregating monomers of PrLD and αS (PrLD–αS hybrid fibrils) or homotypic PrLD fibrils? Similarly, it will be interesting to understand how PrLD–αS hybrid fibrils are organized; are the cross β-sheets are interdigitated β-strands of PrLD and αS? or are they formed as block polymers by epitaxial growth of the two fibrils? In-depth investigations in the future will enable better insights into the structures of PrLD fibril polymorphs, which may uncover molecular basis of phenotype emergence in co-morbid neurodegenerative diseases.

## METHODS

### Recombinant expression and purification of unlabeled PrLD, αS, ^13^C labeled PrLD and ^15^N labeled αS

Recombinant expression and purification of both unlabeled and ^13^C/^15^N labeled TDP-43 PrLD and αS was carried out as described previously (52). For TDP-43 PrLD, N-terminal hexahistidine tag fused construct (Addgene plasmid #98669) with a tobacco etch virus (TEV) cleavage site was expressed in BL21 Star™ (DE3) cells and purified using Ni-NTA affinity chromatography. Briefly, cells expressing unlabeled and labeled TDP-43 PrLD were resuspended in lysis buffer (20 mM Tris, 500 mM NaCl, 5 mM imidazole, 6M urea, 0.5 mM PMSF, pH 8.0), sonicated and centrifuged. The supernatant was incubated with the Ni-NTA beads and washed using wash buffers each containing 15 mM and 30 mM imidazole in buffer containing 20 mM Tris, 500 mM NaCl, 6M urea, pH 8.0 to remove non-specifically bound proteins. Protein was eluted in 150 mM imidazole containing elution buffer followed by dialysis in 20 mM Tris buffer, 500 mM NaCl, 2M urea, pH 8.0, concentrated and stored at −80 °C. For experiments, protein was thawed in ice and desalted using Sephadex G-25 HiTrap desalting column (Cytiva) in 20 mM MES buffer pH 6.0. Protein concentration was determined using UV spectrometer (extinction coefficient 19,480 M^-1^cm^-1^). Both unlabeled and ^15^N labeled recombinant αS was expressed as N-terminus hexahistidine tag followed by thrombin cleavage site in BL21 (DE3) cells. Protein was purified using Ni-NTA affinity chromatography, eluted in elution buffer containing 250 mM imidazole, dialyzed in nanopore water to remove imidazole, concentrated using 10 kDa centrifugal filter (Thermo fisher) and stored at −80 °C. Prior to the experiments, protein was thawed in ice and incubated for 30 minutes with 40 mM NaOH to disaggregate the oligomers and subjected to size exclusion chromatography (SEC) using Superdex-200 column in 20 mM MES buffer pH 6.0. Protein concentration was determined using UV spectrometer using extinction coefficient 5960 M^-1^cm^-1^.

### Labeling of monomeric αS

For cell culture studies, αS was fluorescently labeled using HiLyte™ Fluor 405 succinimidyl ester (Anaspec). Monomeric αS in 20 mM MES buffer pH 6.0 was incubated with 3 molar excess of dye and incubated at 4 °C for 16 h. Excess dye was removed using Sephadex G-25 HiTrap desalting column (Cytiva) for two times and protein was eluted in sterile 20 mM Tris buffer pH 7.5. The protein was used as such for internalization assays.

### Cell growth, transfection and colocalization analysis

Colocalization analysis of αS and TDP-43 PrLD was carried out in SH-SY5Y neuroblastoma cells (ATCC) maintained at 37 °C and 5.5% CO_2_ in DMEM and F-12 (1:1) media containing 10% FBS and 1% Penicillin/Streptomycin (Thermo Fisher Scientific). At first, transfection reagent was prepared by mixing 9 μL of Opti-MEM, 1 μg of TDP-43 PrLD plasmid (sBFP2-PrLD) and TransIT-X2® dynamic delivery system, Mirius (300 μL). The mixture was incubated for 20 minutes prior to transfection in cells with confluency greater than 70% that are grown by seeding 45,000 cells/well in 96 well glass bottom plates (Cellvis). After 24 hours, media containing transfection reagent was replaced, washed for two times with warm fresh media and incubated with approximately 500 nM Hilyte-532 labeled recombinant αS monomers. Cells were further incubated for 24 hours and media containing αS monomers was replaced with fresh media. Cells were stained with nuclear marker (NucSpot® Live 650, Biotium) and imaged using Leica STELLARIS-DMI8 microscope at 40X magnification. All the acquired images were processed using Adobe illustrator software.

### Preparation of homotypic and heterotypic fibrils

Both labeled and unlabeled fibrils samples were prepared for different biophysical studies. For ssNMR, homotypic fibrils of ^13^C labeled TDP-43 PrLD, hybrid fibrils containing ^13^C TDP-43 PrLD and ^15^N αS, and αS^f^-seeded ^13^C labeled TDP-43 PrLD fibrils were used. Homotypic ^13^C labeled TDP-43 PrLD fibrils were prepared by incubating 20 μM ^13^C labeled TDP-43 PrLD monomers for 7 days. Hybrid fibrils were prepared by mixing 20 μM ^13^C TDP-43 PrLD and ^15^N αS at 1:1 stoichiometry and incubating for 7 days. αS^f^-seeded TDP-43 PrLD fibrils were prepared by incubating 2 μM αS sonicated fibrils with 20 μM ^13^C TDP-43 PrLD for 7 days. All the samples were incubated under quiescent conditions in 20 mM MES buffer, 0.01% sodium azide, pH 6.0 at 37 °C. Fibrils from respective reaction were harvested by centrifuging at 20,000 xg for 20 minutes, washed, and centrifuged fibrils pellets were subjected for ssNMR studies. For other studies, unlabeled fibrils were prepared and resuspended in sterile water or buffer prior to the experiment. αS fibrils were prepared by incubating 5 mg of αS monomers with 150 mM NaCl and 0.01% sodium azide in 20 mM Tris buffer pH 8.0 at 37 °C and 600 rpm for 10-14 days. The fibrils were centrifuged at 18,000 xg for 20 minutes, washed, and resuspended in water or buffer of choice. αS sonicated fibrils were generated by sonicating resuspended fibrils using Misonix XL-2000 sonicator for 7 cycles each with 30 seconds burst and one-minute rest at 35% intensity, incubated on ice for 30 minutes prior to incubation with TDP-43 PrLD.

### Au-nanoparticle labeling and transmission electron microscopy (TEM)

All the unlabeled fibrils were prepared as described earlier. Au-nanoparticle-labeled PrLD fibrils were prepared by mixing 5 μM his-tagged PrLD monomers with 20 μM untagged PrLD monomers and incubated under quiescent condition at 37 °C. Moreover, nanogold-labeled hybrid fibrils were prepared by incubating 20 μM untagged PrLD monomers with 5 μM his-tagged αS monomers and 15 μM untagged αS monomers under quiescent condition at 37 °C. Fibrils were harvested by centrifugation and resuspended in ultrapure water. These were directly incubated with 5 nm Ni-NTA-Nanogold® (Nanoprobes Inc,) stock solution at 1:20 v/v dilution. Freshly prepared nanogold-labeled and unlabeled fibrils stock solution was diluted to 1 μM in ultrapure water prior to TEM imaging. All samples were then applied onto ultrathin carbon film supported by a lacey carbon film on 400 mesh copper grids (Ted Pella Inc.) for 2 minutes. Nanogold-labeled samples had an additional wash step of spotting 5 mL of ultrapure water onto the grid for 1 minute two times. Samples were then negatively stained with 2% uranyl acetate for 1 minute. TEM grids were analyzed using a Hitachi HT-7700 TEM.

### Intrinsic tryptophan fluorescence and dye binding assays

Intrinsic tryptophan fluorescence and dye binding of monomers, homotypic and heterotypic fibrils were carried out in Cary Eclipse spectrometer (Varian Inc.) in scan mode. Stock solution of ANS was prepared in water, whereas DCVJ and curcumin were prepared in 95% ethanol. In intrinsic tryptophan fluorescence, samples were equilibrated for 1 min and spectra were collected by setting the excitation and emission wavelength at 280 nm and 300-450 nm, respectively. For ANS, samples were diluted in 100 μM ANS, equilibrated for 1 min and spectra were collected by setting the excitation and emission wavelength of 388 nm and 400-650 nm, respectively. For DCVJ, samples were diluted in 10 μM DCVJ, equilibrated for 1 min before measuring the fluorescence at excitation and emission wavelength of 433 nm and 450-650 nm, respectively. Similarly, curcumin fluorescence was measured by incubating the samples in 5 μM curcumin followed by 1 min equilibration. Fluorescence spectra were collected in excitation and emission wavelength of 430 nm and 470-600 nm, respectively. All the samples were used at 2 μM final concentration in 20 mM MES buffer pH 6.0. The shift in fluorescence emission wavelength maxima was calculated by subtracting the emission wavelength maxima of samples to TDP-43 PrLD monomers. Moreover, percentage change in intensity was calculated by subtracting the integrated area under the curve (AUC) of samples to TDP-43 PrLD monomers.

### Proteinase K digestion

Homotypic and heterotypic fibrils (~ 4 μg) formed by αS and TDP-43 PrLD were resuspended in sterile water and subjected to digestion using 280 ng of proteinase K (Ambion, Inc.) diluted from a stock of 20 mg/mL in sterile water. All the reactions were carried out by shaking at 200 xg at 37 °C. Reactions were quenched at 10, 20, 30, and 40 minutes, respectively using 0.5 mM PMSF. Finally, aliquots of reactions were mixed with non-reducing laemmli sample buffer (4X) and subjected to SDS-PAGE. Gel was stained using Pierce™ silver stain kit (ThermoFisher scientific) as per manufacturer’s protocol and imaged on a GelDoc molecular imager (Bio-Rad).

### MALDI-ToF spectrometry

PK digestion of monomers, homotypic and heterotypic fibrils (1 μg) was carried out as described before. Samples for MALDI-ToF were prepared by mixing 1 μL of the digested fibrils with equal volume of sinapinic acid prepared in 1:1 acetonitrile/water and 0.1% TFA. The sample was spotted on the MSP 96 MALDI plate (Bruker Daltonics). Spectra were collected by maintaining the laser power at 70% and analyzed using flex analysis. All the data were processed using OriginPro 8.5.

### Circular dichroism (CD) melting

SDS-induced temperature melt of TDP-43 PrLD monomers or homotypic fibrils, hybrid fibrils, and αS^f^-seeded PrLD fibrils was carried out on a Jasco J-815 spectrometer. The fibrils were resuspended in water and spectra were collected in 10 mm pathlength cuvette at standard sensitivity with scan rate of 50 nm/min, 8 seconds DIT, 1 nm bandwidth, and 0.1 nm data pitch. For stability analysis, 1% w/v SDS was added to the samples, immediately followed by temperature interval measurement by collecting the spectra at every 5 °C temperature interval from 20 °C to 90 °C. The data were normalized for mean residual ellipticity (MRE) and ellipticity difference between 208 nm and 218 nm was calculated. The ellipticity difference, as an indicative of α-helical or β-sheet structure, was plotted against the temperature, and fit using the Boltzman function in Origin 8.5.

### Solid-state NMR

Fibrils were packed into Bruker 3.2 mm NMR rotors via ultracentrifugation at 4 °C and 150, 000 x *g* for 30 minutes using Beckman Optima XPN-100 fitted with a SW-41 Ti swinging-bucket rotor and Ultra-Clear tubes. NMR spectra were collected on a 11.75 T magnet (500 MHz ^1^H NMR frequency) using a Bruker spectrometer and a 3.2mm Bruker Low-E ^1^H/^13^C/^15^N NMR probe. Two-dimensional (2D) ^13^C-^13^C NMR spectra were collected using the 2D dipolar assisted rotational resonance (DARR) NMR technique at 10 kHz magic angle spinning speed with a mixing time of 50 ms. A ^1^H radiofrequency field of 100 kHz was used. Signal averaging of all the spectra in Figure 6 required approximately 24 h. This technique produces off-diagonal peaks (crosspeaks) that mostly correspond to interactions between ^13^C atoms within the same amino acid (63, 84, 85).

### Primary neuron isolation and cell treatment

This study was conducted in a facility approved by the American Association for the Accreditation of Laboratory Animal Care. All procedures were performed in accordance with recommendations in the Guide for the Care and Use of Laboratory Animals of the National Institutes of Health. Our protocol was approved by the Institutional Animal care and Use Committee of the University of Texas Medical Branch (UTMB). Primary cortical neuronal cultures were prepared and maintained as described previously (61, 86–88). Briefly, cortices were isolated from C57BL/6 mice (Jackson Laboratory; 000664) during embryonic day 13-16. Brain tissues were digested using Accutase solution (Sigma) followed by gentle trituration with a fire-polished glass pasture pipet. Dissociated cells was plated at a density of 1.6 x 10^5^ cells/ml on poly-L-lysine coated coverslips. Culture media contains neurobasal medium (Gibco; 12348017) supplemented with 2% B-27 Plus supplement (Gibco; A3582801), 0.5 mM GlutaMax (Gibco; 35050-061), 10,000 units/ml penicillin, 10,000 μg/ml streptomycin, and 25 μg/ml amphotericin B (Gibco; 15240062). Half of the media was changed every 3-4 days. On 10 days *in vitro,* cells were treated with monomers or fibrils of α-synuclein, TDP-43 PrLD, or αS^f^-seeded PrLD fibrils at 1-2 μM for 24 h. Buffers used in samples preparation were the vehicle control in all experiment.

### Immunofluorescence, confocal microscopy, and imaging analysis

After cell treatments, primary neurons were gently washed 3 times with 1X PBS. Formaldehyde solution 4% (Sigma) was used for fixation for 15 min at RT followed by 3 washes. Cells were permeabilized using 0.25% Triton X-100 (Sigma) in 1X PBS for 10 min and blocked for 30 min at RT in blocking buffer. Cells were then incubated with primary antibodies diluted in blocking buffer overnight at 4°C. Primary antibodies used include mouse anti-βIII-tubulin (1:1000, Abcam; ab78078), rabbit anti-Synapsin I (1:1000, Abcam; ab8), or rabbit anti-PSD95 (1:1000, Abcam; ab18258). On the next day, cells were washed and incubated with secondary antibodies (1:700, Life Technologies) for 1 h at RT in the dark. After three washes, cells were mounted with Prolong Gold antifade reagent with DAPI. All samples were examined with 63x objective of a Zeiss LSM 880 confocal microscope using 405-nm diode laser and argon laser 458/488/514 nm. To build the z-stack, 17 stacks/0.37–0.41-μm optimal thickness were captured. Each treatment condition was randomly imaged in five different regions of interest and performed in duplicate. Neuron-specific pre-and post-synaptic signals were analyzed using Imaris software as previously described (89). Briefly, for each image, pre- and post-synaptic channels were filtered based on neuronal surface and discrete quantitative “spots” were created. Once pre- and post-synaptic spots were detected, the distance threshold to identify colocalized spots was set to 1mm, providing for an estimate of synapse number (90–92). The number of co-localized spots were normalized to the untreated. Statistical analysis was performed in Graph Pad Prism 9 using One-way ANOVA followed by Tukey’s Test. Results were considered statistically significant at *p* < 0.05.

## Acknowledgements

The authors would like to thank the following agencies for financial support: National Institute of Aging (1R56AG062292-01 (to VR); AG054025, RF1AG077484 (to RK) and RF1AG073434-01A1 (to AP), the National Institute on Minority Health and Health Disparities (RF1AG073434-01A1; to AP), and the National Science Foundation (NSF CBET 1802793) to VR.

The authors also thank the National Center for Research Resources (5P20RR01647-11) and the National Institute of General Medical Sciences (8 P20 GM103476-11) from the National Institutes of Health for funding through INBRE for the use of their core facilities, and the National Science Foundation (NSF MRI 2019023) for STED confocal microscope facility. The authors acknowledge NMR facility at Georgia Tech and Robert P. Apkarian Integrated Electron Microscopy Core at Emory. The authors would also like to thank Dr. Jonathan Lindner for his gracious help with the confocal microscope use and troubleshooting.

## Author contributions

VR conceptualized the project and was involved in all aspects of the manuscript especially biophysical studies; AP contributed significantly to idea development and in manuscript preparation along with directing the NMR and TEM work. RK directed primary neuron studies and provide intellectual support. SD performed protein expression and purification, sample preparations, all biophysical experiments, co-localization cell culture experiments along with intellectual discussions and manuscript writing; AR performed NMR and TEM imaging experiments and in manuscript editing; NB, NP and LF performed primary neuronal cell culture experiments along with data and image processing.

